# Lifespan in rodents with MYT1L heterozygous mutation

**DOI:** 10.1101/2024.10.30.621094

**Authors:** Allyson Schreiber, Raylynn G. Swift, Leslie Wilson, Kristen L. Kroll, Joseph D. Dougherty, Susan E. Maloney

## Abstract

MYT1L syndrome is a newly recognized disorder characterized by intellectual disability, speech and motor delay, neuroendocrine disruptions, ADHD, and autism. In order to study this gene and its association with these phenotypes, our lab recently created a *Myt1l* heterozygous mutant mouse inspired by a clinically relevant mutation. This model recapitulates several of the physical and neurologic abnormalities seen in humans with MYT1L syndrome, such as weight gain, microcephaly, and behavioral disruptions. The majority of patients with this syndrome are young, and little is known about the impact of age on health and mortality in these patients. Using a *Myt1l* mutant mouse, we examined the impact of *Myt1l* mutation on body weights, lifespan, and histopathology findings of mice at the end of life. This cohort of heterozygous mice demonstrated increased body weight across the lifespan, however there was no significant difference in lifespan, apparent cause of death, or end of life histopathological findings between *Myt1l* heterozygous and wildtype mice. These findings suggest while *Myt1l* heterozygous mutation may influence overall brain development, it does not strongly impact other organ systems in the body over time.

## Background

Neurodevelopmental disorders, including autism spectrum disorder, intellectual disability, and attention deficit hyperactivity disorder, affect more than 3% of children worldwide and lead to impaired cognition, communication, adaptive behavior, and psychomotor skills (1,2). Multiple genetic syndromes have been associated with neurodevelopmental disorders including autism. People with neurodevelopmental disorders often have shorter lifespans than the general population, likely due to a variety of factors including comorbid conditions and health care disparities (3,4).

Recently, the gene Myelin Transcription Factor 1 Like (*MYT1L*) has been associated with neurodevelopmental disorders (NDD), with *MYT1L* loss of function now recognized as MYT1L Syndrome (1). Hallmark features of MYT1L Syndrome include intellectual disability, obesity, speech and motor delay, neuroendocrine disruptions, ADHD and autism. Epilepsy, microcephaly, and white matter thinning are also observed in a portion of patients (5–9). While significant progress has been made in characterizing the molecular and cellular mechanisms that underlie Myt1L syndrome (10–13), the majority of identified patients are young (<35 years), and there is still much to learn about the long-term impact of *Myt1L* gene mutations on overall health outcome. Syndromic NDDs can sometimes include comorbidities not apparent until later in life, such as elevated risk of particular cancers in Tatton-Brown-Rahmen Syndrome (TBRS) or early onset Alzheimer’s disease in Down Syndrome (14,15). Notably, it is unknown if *MYT1L* mutation may result in any such recurrent comorbidities that would influence overall lifespan and cause of death.

While waiting for definitive studies in humans, study of lifespan in animal models can be helpful to understand potential long-term health impacts of newly discovered genetic mutations. Mice have substantially shorter lifespans than humans, enabling studies of how a particular genetic mutation intersects with time to impact health. Most mouse strains are generally considered geriatric at approximately 24 months (16); however, lifespan is reported to vary between strains, with C57BL/6 mice known to be long-lived with 50% survival at approximately 900 days of life (17). At least three mouse models of syndromic NDD show decreased lifespan including Down Syndrome, Prader-Willi, and Rett Syndrome (15,18–22), while others like TBRS either do not impact lifespan or have not yet been studied (14).

We previously generated and characterized a mouse mutant disrupting the *Myt1l* gene (10). These studies demonstrated that *Myt1l* mutant mice exhibit a range of neurological and physical abnormalities, including altered neuronal function, behavior, and body weight regulation. The impact of *Myt1l* gene mutations on lifespan and cause of death in these mice has not yet been explored. Studying the lifespan and cause of death in *Myt1l* mutant mice will be helpful in understanding the potential health implications of *MYT1L* gene mutations in humans. In this paper, we examine differences in lifespan, gross necropsy, and histopathological findings between *Myt1l* heterozygous mutant and wildtype mice at the end of life.

## Methods

### Animals

All experimental protocols were approved by and performed in accordance with the relevant guidelines and regulations of the Institutional Animal Care and Use Committee of Washington University in St. Louis and were in compliance with US National Research Council’s Guide for the Care and Use of Laboratory Animals, the US Public Health Service’s Policy on Humane Care and Use of Laboratory Animals, and Guide for the Care and Use of Laboratory Animals. This study is reported in accordance with ARRIVE guidelines.

All mice used in this study were maintained and bred in the vivarium at Washington University in St. Louis School of Medicine. The colony room lighting was on a 12:12 h light/dark cycle (lights on at 6a.m.); room temperature (20–22C) and relative humidity (50%) were controlled automatically. Standard lab diet and water were available *ad lib*. Upon weaning at postnatal day (P)21, mice were group housed according to sex and experimental condition. The mice used in this study harbor a frameshift mutation in exon 11 of the *Myt1l* gene on a C57BL/6J background, as previously described (10). The cohort used herein consisted of 16 *Myt1l* heterozygous mutants (‘Het’, 8 males, 8 females) and 21 wildtype littermate controls (‘WT’, 8 males, 13 females). All mice reported here were used for behavioral testing and magnetic resonance imaging between P33 and P287, as published in Chen et al (2021). A subset of animals including five Hets (two males, three females) and six WT (two males, four females) were submitted for gross necropsy and histopathological examination. Following completion of the study, mice were humanely euthanized with carbon dioxide overdose in accordance with the American Veterinary Medical Association guidelines.

### Moribund Status Determination

General health status and body weight were monitored on a weekly basis from P383 until P720. Monitoring continued for moribund state or mortality until P1013-1015. Moribund mice were euthanized via carbon dioxide overdose if judged to be severely ill and/or exhibiting signs such as gulping or irregular breathing; severe motor/gait disturbance (lack of spontaneous movement and little to no movement when prompted); ulcerated skin, or abdominal distension. The date of euthanasia was used as an estimate of natural lifespan in these cases. The experiment was continued until day 1013-1015, and mice that had survived to that point were considered censored (not plotted in figures).

### Histopathology

Gross necropsy, tissue processing, and slide staining were performed by the Research Animal Diagnostic Laboratory at Washington University in St. Louis School of Medicine. Tissues collected at the time of gross necropsy were fixed in 10% neutral buffered formalin for 24-48 hours, paraffin embedded, sectioned at 5-µm thickness and stained with hematoxylin and eosin (H&E, Harris Hematoxylin Nuclear Stains, Cat. No. 3801560). Histopathological evaluation was performed by a board-certified veterinary pathologist. Animals examined included five *Myt1l* Hets (two males, three females) and six wild types (two males, four females).

### Statistical Analysis

Statistical analyses and data visualization were conducted using IBM SPSS Statistics (v.28). Prior to analyses, weight data was screened for missing values and fit of distributions with assumptions underlying univariate analysis. This included the Shapiro-Wilk test on z-score-transformed data and qq-plot investigations for normality, Levene’s test for homogeneity of variance, and boxplot and z-score (±3.29) investigation for identification of influential outliers. Analysis of variance (ANOVA) was used to analyze weight data, and simple main effects were used to dissect significant interactions. Kaplan–Meier survival analysis was conducted to assess lifespan. Sex was included as a biological variable in all analyses across all experiments. Multiple pairwise comparisons were subjected to Bonferroni correction. The critical alpha value for all analyses was p < .05. Figure illustrations were generated using Prism software. The datasets generated and analyzed during the current study are available from the corresponding author upon reasonable request.

## Results

At the end of our initial behavioral and neuroimaging-based characterization of *Myt1l* mutants (10), we continued housing the animals until they became moribund. This allowed us to examine the health and lifespan of a cohort of Het and WT littermates, over two to three years, with a subset further assessed grossly and histologically at time of death.

### *MYT1L* heterozygous mutant mice weigh more than wild type mice into old age

Previously, we observed a significant increase in body weight starting in early adulthood in mice harboring a *Myt1l* mutation (Fig 1A; Chen et al. 2021). Here, we have extended the analysis of body weight into old age (to P720) to determine if *Myt1l* mutation effects on body weight persisted. We ran a three-way ANOVA to examine the effect of sex, genotype, and age on weight data collected weekly between approximately P530 and P720 (Fig 1B). There was no significant three-way interaction, *F*(21,671) = 48.03, *p* = 1.00, but significant main effects of sex (*F*(1,671) = 57.35, *p* = 0.000), genotype (*F*(1,671)=80.16, *p*=0.000) and a significant sex*genotype interaction (*F*(1,671) = 32.15, *p* = 0.000) were found. There was no significant main effect of age or significant interactions with age. Female Het mice (*M* = 36.98, *SE* = 0.52) were significantly heavier than female WT mice (*M* = 30.50, *SE* = 0.35), *F*(1,671)= 106.40; *p* =0.00. Male Het mice (*M* = 37.83, *SE* = 0.44) were significantly heavier than male WT mice (*M* =36.37, *SE* = 0.44), *F*(1,671) = 5.42 , *p* = .02. Expected sex differences were found in WT animals, with males heavier than females, *F*(1,671) = 107.6, *p* = 0.00., but there was no significant difference in weight between male and female Het mice, *F*(1,671) = 1.52; p=0.136.

**Figure 1:**
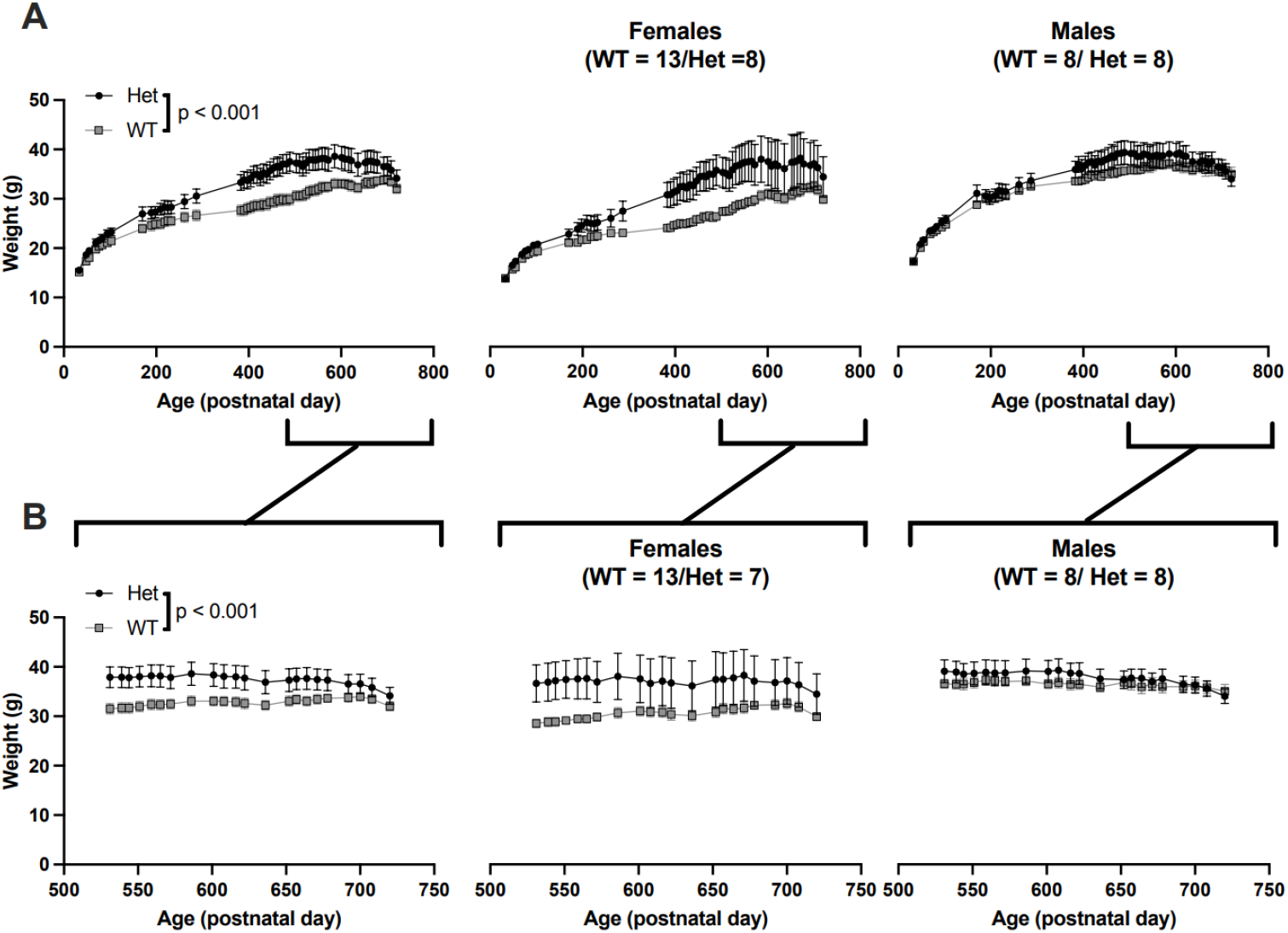
Het mice weighed more than WT throughout the lifespan. A) Weights of cohort across age, including data previously reported in Chen et al (prior to P500), and newly collected data. Left panel, all mice, right panels, subsetted by sex. B) As above, plotting only newly collected data,3 way ANOVA, for age sex and genotype, main effect of genotype shown.

### *MYT1L* heterozygous mutation does not impact lifespan in mice

To understand if heterozygous mutation for *MYT1L* influences lifespan, we continuously monitored the status of our mice into their old age. We performed a Kaplan–Meier survival analysis over the lifespan of male and female Hets and WT littermates. Date and cause of death were noted for all mice. At ∼P1014 or >33 months, all surviving animals were euthanized, which included 4 Het males, 1 WT male and 3 WT females. We found a significant difference in survivability between males and females (*χ*= 9.61, p = 0.002; Fig 2A). Specifically, males in our cohort lived longer than females. Males, pooled across genotypes, had a longer median lifespan (958.5 days) than females (777 days). However, we did not observe a significant difference between Het and WT animals (*χ* = 0.95, p = 0.330; Fig 2B), with WT animals achieving a median lifespan of 875 days compared to 762.5 days for Hets. Due to small group sizes (<20/group), genotype*sex interactions were not analyzed or interpreted.

**Figure 2:**
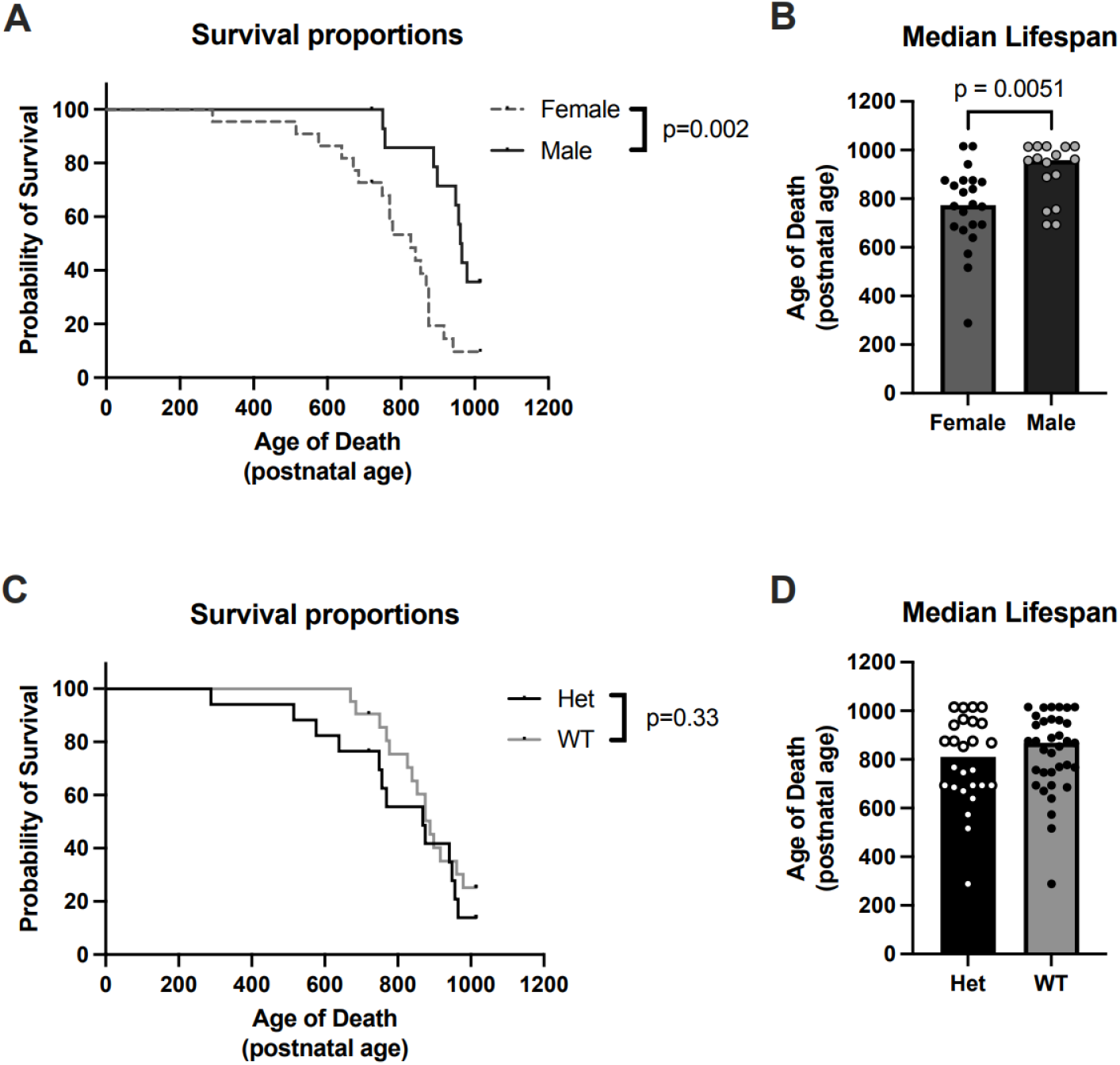
Het and WT mice have similar lifespans. A) A Kaplan-Meier plot of survival comparing male and female collapsed for genotype mice show longer lifespans in males. B) Bar chart illustrating median lifespan for age of death, including animals at end of experiment (p1013-1015). Filled circles are individual animals. C) A Kaplan-Meier plot of survival comparing Het and WT mice shows no significant genotype difference in lifespan. D) Bar chart illustrating median lifespan for age of death across genotype, including animals at end of experiment (p1013-1015). Filled circles are individual animals.

### *Myt1l* heterozygous mutation does not result in significantly different pathology at death

To understand if *Myt1l* Het mice experienced similar health outcomes with old age as compared to their WT littermates, a subset of animals were submitted for gross necropsy and histopathological examination once they were judged to be moribund or found dead. The subset of mice that were examined for gross necropsy and histopathology were on average 820.9 days old (2.2 years old) for WT mice and 773.5 days old (2.1 years old) for Het mice, both groups well into old age and not statistically different from each other (p=0.34) (16,23). We found that mice harboring a *Myt1l* mutation had similar gross and histopathological findings as compared to WT littermate controls (Tables 1 and 2). Specifically, we found similar age-related lesions including cancers and changes in liver, kidney and bone marrow morphologies. Cancers, such as lymphoma, leukemia, and hepatocellular carcinoma were identified in most animals of both genotypes (4/5 WT animals, 3/5 Het animals). Extramedullary hematopoiesis, the production of red and white blood cells outside of bone marrow, was found in both groups (2/5 WT animals, 5/5 Het animals). Underlying causes of extramedullary hematopoiesis include anemia, chronic inflammation, and neoplasia, including lymphoma and leukemia. Membranoproliferative glomerulopathy, a kidney disorder that ultimately affects the kidney’s ability to adequately filter blood and create urine, was found in both WT (3/5) and Het animals (3/5). Liver changes, such as oval cell and Kupffer cell hyperplasia were also observed (both 2/5). Biliary cyastadenoma was found in two WT animals. This benign liver malformation is uncommonly described in mice and best characterized as a bile duct hamartoma (von Meyernburg complex). In these two animals, the masses were large enough to cause abdominal distension with compression of other internal organs, resulting in rectal prolapse in one mouse. Following statistical analysis, we did not determine there to be an increased incidence in specific organ changes or disease processes in *Myt1l* Het mice as compared to WT littermates.

**TABLE 1:**
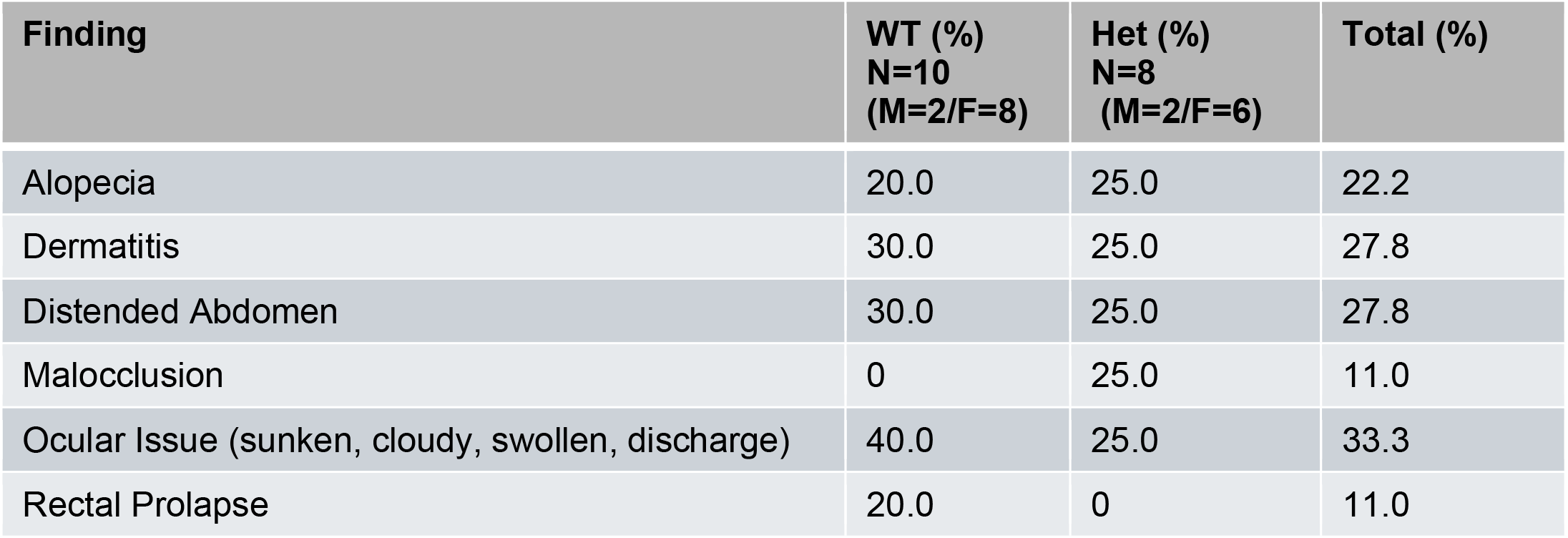
Gross Necropsy Findings.

**TABLE 2:**
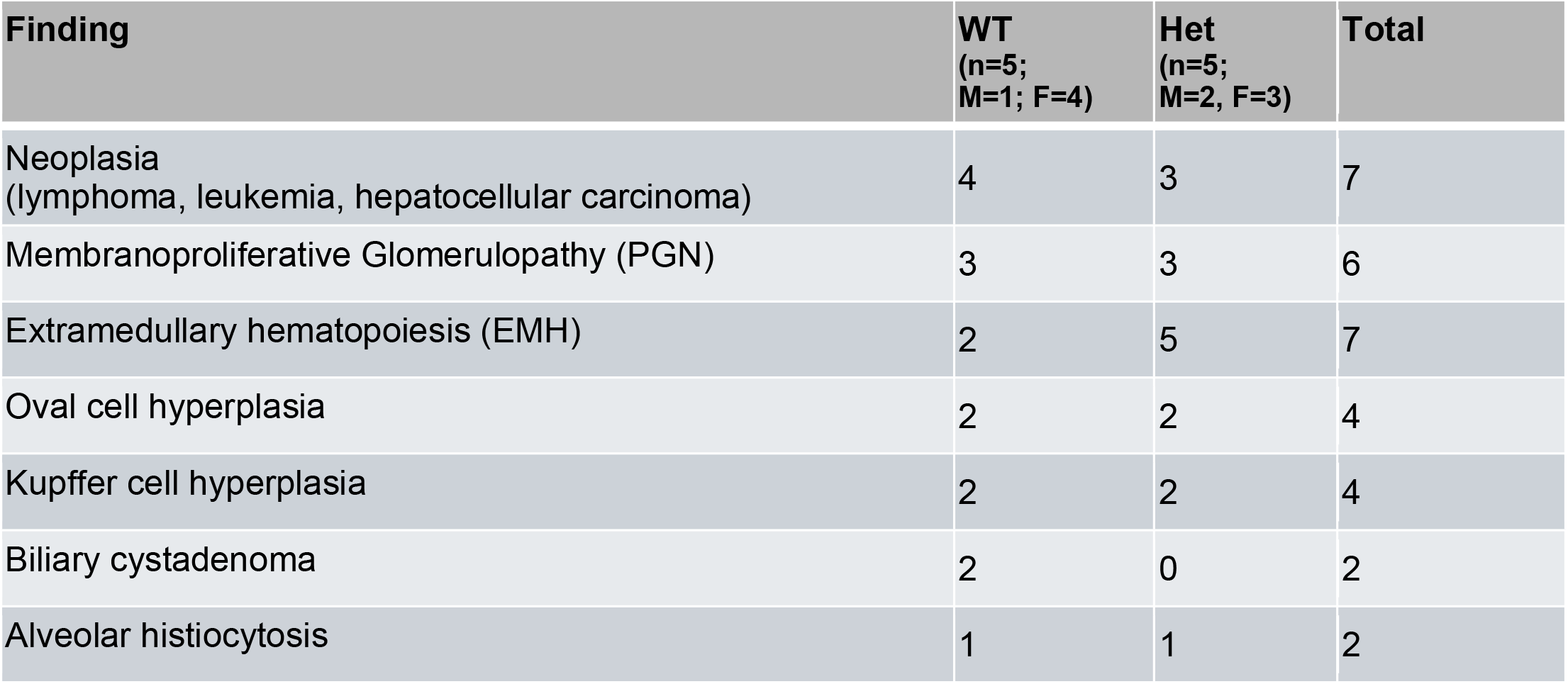
Histopathology Findings.

## Discussion

MYT1L Syndrome is a newly defined monogenic form of NDD, and by studying recently generated mutant mouse models, we are beginning to understand how *MYT1L* mutations alter brain development and contribute to NDD-related features. Using mouse models of disease, we can study pathologies across lifespan and into old age, in hopes of identifying potential clinical complications and comorbities in human patients as they age. In this paper, we examined the lifespan of a cohort of *Myt1l* mutant mice and cataloged gross and histological changes to understand possible end of life consequences of *Myt1l* mutation. This cohort of *Myt1l* heterozygous mice continued to have increased body weight into old age compared to WT counterparts but did not have consistent differences in lifespan or necropsy findings.

Approximately 50% of people with MYT1L syndrome exhibit overweight/obesity, potentially due to hyperphagia or neuroendocrine disturbances (24). Previously we have shown that this cohort of adult heterozygous MYT1L mice had higher body weights, compared to wildtype controls (10), which was maintained through the duration of this study. However, we do note that an obesity-related increase in weight was not reliably found in all future cohorts (Maloney et al, *in prep*). Although there was no significant effect of age on weight in this aging cohort, there was greater variability in weight scores in Het mice than WT mice, especially at the later timepoints and heavily driven by females. Finally, although not examined specifically in detail, there did not appear to be any increased weight-related changes related to cause of death at time of histopathological analysis.

We did not see a significant difference in survival between *Myt1l* Het mice and WT mice, whereas mortality studies in other animal models of NDD genetic liability have demonstrated mixed results. For example, genetically engineered mouse models of Prader-Willi syndrome have shown significant variation in lifespan attributed primarily to neonatal mortality, which is dependent on the type of genetic model used (22). In addition, while several different genetic rodent models of Rett syndrome have shown comparable decreases in lifespan, these models have also demonstrated variability in cause of death (respiratory failure versus kidney failure), depending on the specific model used (20,21,25). With additional, novel discoveries of genetic causes of NDD, such as MYT1L syndrome, studies on aging and lifespan will hopefully provide additional insight into the pathophysiology of aging in individuals with this disorder, in hopes of identifying potential causes of comorbidities and death. However, these studies are currently limited or still in progress.

Despite no group effects on survival between *Myt1l* Het mice and WT mice, there was a significant sex difference on overall lifespan, with male mice living significantly longer than female mice. Sex differences on lifespan of various inbred mouse strains, including the C57Bl/J have been reported, but have been mixed across studies and institutions, with some showing males outlive females, others showing females outlive males, and some showing no difference in lifespan at all (26). Thus, the sex difference found in our survival analysis could be an effect of background strain or the influence of a number of facility-specific and/or cohort-specific factors, such as the handling and behavioral testing experience of these animals.

We also examined histopathology in a subset of animals. *Myt1l* Het mice had similar histopathological findings as their WT littermate controls. Common findings in the histological examination were likely related to old age including benign and malignant neoplasms, membranoproliferative glomerulonephropathy, extramedullary hematopoiesis, and aging changes in the liver. In human studies, people with MYT1L syndrome primarily have central nervous system and endocrine-related pathologies, which we did not specifically see in our histopathological analysis (24). Unlike other syndromic neurodevelopmental disorders, such as Rett syndrome or Prader-Willi where the cause of the death seems to be related to organ dysfunction, there is still not enough information in patients with MYT1L syndrome to understand if specific organs are adversely affected (22,25). In this study, we did not see differences in histopathological changes between *Myt1l* Het mice and WT controls, suggesting old age had a bigger impact on organ function than loss of *Myt1l*. However, more natural history studies in humans and aging studies in mice are needed to definitively rule out potential pathological organ changes in MYT1L syndrome.

The purpose of this study was to understand if the presence of Myt1l mutation impacts lifespan. While mortality in old age may have been similar between Het mice and their wildtype littermates, this study did not target potential sub-lethal disease states throughout life. If there were earlier onset comorbidities, overall age-related changes at >24 months could have masked differences between the two groups. Thus, it remains uncertain whether any pathologies could have presented at an earlier age, as this study provided only a snapshot of health and disease at the end of life. A similar, future study with a larger cohort of mice, to include groups sampled for necropsy and histopathology at early adulthood (2-3 months), mid-life (10-14 months) and early onset of old age (18 months) could provide greater insight into potential sub-lethal disease states or other pathophysiologies associated with MYT1L syndrome in humans. In addition to examining a wider breadth of ages, it would also be worthwhile to examine specific organs and/or cell types, in hopes of possibly elucidating more subtle changes potentially contributing to the overall morbidity of MYT1L heterozygotes that were not examined in this survey study. If these differences do exist, they did not seem to influence overall lifespan, in the current study, however.

Finally, this study included only a small number of animals tested on a single background strain (C57BL/6J), which is consistently reported as especially long-lived among inbred mouse strains (17,27), with a max lifespan estimated at 1075 ± 13 days in females and 1061 ± 17 days in males (17). As there are known differences in disease development and progression between mouse strains, as well as documented species differences between mice and humans, it remains possible that increased morbidity and/or mortality in humans with MYT1L syndrome might not be detected in this particular mouse model. Causes of decreased lifespan amongst people with neurodevelopmental disorders and autism are presumed to be multifactorial, including influencers such as social determinants of health and access to healthcare, which are not recapitulated in animal models (3,4). Therefore, although there were no definitive differences in lifespan or cause of death between our *Myt1l* Het mice and WT controls, this does not preclude potential lifespan differences in humans with MYT1L syndrome. As MYT1L syndrome becomes recognized and diagnosed with increasing frequency, future studies in animals and humans will be essential for understanding both the lifespan and the healthspan as these patients age. Nonetheless, the current findings suggest a robust lifespan in the context of MYT1L mutation is possible.

## Acknowledgements

We would like to thank members of the Dougherty lab for edits and advice. This work was supported by R01MH124808 (to KLK, JDD, SEM) and the Intellectual and Developmental Disabilities Research Center (IDDRC@WUSTL, P50HD103525)

## Declarations

### Ethical Approval

All experimental protocols were approved by and performed in accordance with the relevant guidelines and regulations of the Institutional Animal Care and Use Committee of Washington University in St. Louis and were in compliance with US National Research Council’s Guide for the Care and Use of Laboratory Animals, the US Public Health Service’s Policy on Humane Care and Use of Laboratory Animals, and Guide for the Care and Use of Laboratory Animals.

### Additional Information Competing Interests

The authors declare no competing interests.

## Data Availability Statement

The datasets generated and analyzed during the current study are available from the corresponding author upon reasonable request.

## Author Contributions

J.D.D. and S.E.M. designed the study. R.G.S. and L.W. collected and analyzed the data. R.G.S., A.S., and S.E.M. wrote the manuscript. A.S., K.L.K, S.E.M, and J.D.D. revised the manuscript and secured funding for the project.

## REFERENCES

1. Boyle CA, Boulet S, Schieve LA, Cohen RA, Blumberg SJ, Yeargin-Allsopp M, et al. Trends in the prevalence of developmental disabilities in US children, 1997-2008. Pediatrics. 2011 Jun;127(6):1034–42.

2. Association AP. Neurodevelopmental Disorders: DSM-5® Selections. American Psychiatric Pub; 2015. 198 p.

3. Perkins EA, Moran JA. Aging Adults With Intellectual Disabilities. JAMA. 2010 Jul 7;304(1):91–2.

4. Hirvikoski T, Mittendorfer-Rutz E, Boman M, Larsson H, Lichtenstein P, Bölte S. Premature mortality in autism spectrum disorder. Br J Psychiatry J Ment Sci. 2016 Mar;208(3):232–8.

5. De Rocker N, Vergult S, Koolen D, Jacobs E, Hoischen A, Zeesman S, et al. Refinement of the critical 2p25.3 deletion region: the role of MYT1L in intellectual disability and obesity. Genet Med Off J Am Coll Med Genet. 2015 Jun;17(6):460–6.

6. Blanchet P, Bebin M, Bruet S, Cooper GM, Thompson ML, Duban-Bedu B, et al. MYT1L mutations cause intellectual disability and variable obesity by dysregulating gene expression and development of the neuroendocrine hypothalamus. PLoS Genet. 2017 Aug;13(8):e1006957.

7. Loid P, Mäkitie R, Costantini A, Viljakainen H, Pekkinen M, Mäkitie O. A novel MYT1L mutation in a patient with severe early-onset obesity and intellectual disability. Am J Med Genet A. 2018 Sep;176(9):1972–5.

8. Santos JFD, Acosta AX, Scheibler GG, Pitanga PML, Alves ES, Meira JGC, et al. Case of 15q26-qter deletion associated with a Prader-Willi phenotype. Eur J Med Genet. 2020 Aug;63(8):103955.

9. Windheuser IC, Becker J, Cremer K, Hundertmark H, Yates LM, Mangold E, et al. Nine newly identified individuals refine the phenotype associated with MYT1L mutations. Am J Med Genet A. 2020 May;182(5):1021–31.

10. Chen J, Lambo ME, Ge X, Dearborn JT, Liu Y, McCullough KB, et al. A MYT1L syndrome mouse model recapitulates patient phenotypes and reveals altered brain development due to disrupted neuronal maturation. Neuron [Internet]. 2021 Oct 5 [cited 2021 Oct 20]; Available from: https://www.sciencedirect.com/science/article/pii/S0896627321006814

11. Chen J, Fuhler NA, Noguchi KK, Dougherty JD. MYT1L is required for suppressing earlier neuronal development programs in the adult mouse brain. Genome Res. 2023 Apr 1;33(4):541–56.

12. Wöhr M, Fong WM, Janas JA, Mall M, Thome C, Vangipuram M, et al. Myt1l haploinsufficiency leads to obesity and multifaceted behavioral alterations in mice. Mol Autism. 2022 May 10;13(1):19.

13. Weigel B, Tegethoff JF, Grieder SD, Lim B, Nagarajan B, Liu YC, et al. MYT1L haploinsufficiency in human neurons and mice causes autism-associated phenotypes that can be reversed by genetic and pharmacologic intervention. Mol Psychiatry. 2023 Feb 14;1–14.

14. Ostrowski PJ, Tatton-Brown K. Tatton-Brown-Rahman Syndrome. In: Adam MP, Feldman J, Mirzaa GM, Pagon RA, Wallace SE, Bean LJ, et al., editors. GeneReviews® [Internet]. Seattle (WA): University of Washington, Seattle; 1993 [cited 2024 Aug 26]. Available from: http://www.ncbi.nlm.nih.gov/books/NBK581652/

15. Ruparelia A, Pearn ML, Mobley WC. Aging and intellectual disability: Insights from mouse models of down syndrome. Dev Disabil Res Rev. 2013;18(1):43–50.

16. Jackson SJ, Andrews N, Ball D, Bellantuono I, Gray J, Hachoumi L, et al. Does age matter? The impact of rodent age on study outcomes. Lab Anim. 2017 Apr 1;51(2):160–9.

17. Yuan R, Peters LL, Paigen B. Mice as a mammalian model for research on the genetics of aging. ILAR J. 2011;52(1):4–15.

18. Bird LM, Billman GF, Lacro RV, Spicer RL, Jariwala LK, Hoyme HE, et al. Sudden death in Williams syndrome: report of ten cases. J Pediatr. 1996 Dec;129(6):926–31.

19. Relkovic D, Doe CM, Humby T, Johnstone KA, Resnick JL, Holland AJ, et al. Behavioural and cognitive abnormalities in an imprinting centre deletion mouse model for Prader–Willi syndrome. Eur J Neurosci. 2010;31(1):156–64.

20. Patterson KC, Hawkins VE, Arps KM, Mulkey DK, Olsen ML. MeCP2 deficiency results in robust Rett-like behavioural and motor deficits in male and female rats. Hum Mol Genet. 2016 Aug 1;25(15):3303–20.

21. Wu Y, Zhong W, Cui N, Johnson CM, Xing H, Zhang S, et al. Characterization of Rett Syndrome-like phenotypes in Mecp2-knockout rats. J Neurodev Disord. 2016;8:23.

22. Zahova S, Isles AR. Chapter 29 - Animal models for Prader–Willi syndrome. In: Swaab DF, Buijs RM, Lucassen PJ, Salehi A, Kreier F, editors. Handbook of Clinical Neurology [Internet]. Elsevier; 2021 [cited 2024 Aug 19]. p. 391–404. (The Human Hypothalamus; vol. 181). Available from: https://www.sciencedirect.com/science/article/pii/B9780128206836000294

23. Flurkey K, M. Currer J, Harrison DE. Chapter 20 - Mouse Models in Aging Research. In: Fox JG, Davisson MT, Quimby FW, Barthold SW, Newcomer CE, Smith AL, editors. The Mouse in Biomedical Research (Second Edition) [Internet]. Burlington: Academic Press; 2007 [cited 2024 Aug 19]. p. 637–72. (American College of Laboratory Animal Medicine). Available from: https://www.sciencedirect.com/science/article/pii/B9780123694546500741

24. Coursimault J, Guerrot AM, Morrow MM, Schramm C, Zamora FM, Shanmugham A, et al. MYT1L-associated neurodevelopmental disorder: description of 40 new cases and literature review of clinical and molecular aspects. Hum Genet. 2022 Jan;141(1):65–80.

25. Ward CS, Huang TW, Herrera JA, Samaco RC, Pitcher MR, Herron A, et al. Loss of MeCP2 Causes Urological Dysfunction and Contributes to Death by Kidney Failure in Mouse Models of Rett Syndrome. PloS One. 2016;11(11):e0165550.

26. Austad SN, Fischer KE. Sex Differences in Lifespan. Cell Metab. 2016 Jun 14;23(6):1022–33.

27. Goodrick CL. Life-span and the inheritance of longevity of inbred mice. J Gerontol. 1975 May;30(3):257– 63.

