## Supplemental Table 1 for "Lifespan in rodents with MYT1L heterozygous mutation"

**TABLE 1: Gross Necropsy Findings**

| **Finding** | **WT (%)**  **N=10 (M=2/F=8)** | **Het (%)**  **N=8**  **(M=2/F=6)** | **Total (%)** |
| --- | --- | --- | --- |
| Alopecia | 20.0 | 25.0 | 22.2 |
| Dermatitis | 30.0 | 25.0 | 27.8 |
| Distended Abdomen | 30.0 | 25.0 | 27.8 |
| Malocclusion | 0 | 25.0 | 11.0 |
| Ocular Issue (sunken, cloudy, swollen, discharge) | 40.0 | 25.0 | 33.3 |
| Rectal Prolapse | 20.0 | 0 | 11.0 |
