## Supplemental Table 2 for "Lifespan in rodents with MYT1L heterozygous mutation"

**TABLE 2: Histopathology Findings**

| **Finding** | **WT**  **(n=5;**  **M=1; F=4)** | **Het**  **(n=5;**  **M=2, F=3)** | **Total** |
| --- | --- | --- | --- |
| Neoplasia  (lymphoma, leukemia, hepatocellular carcinoma) | 4 | 3 | 7 |
| Membranoproliferative Glomerulopathy (PGN) | 3 | 3 | 6 |
| Extramedullary hematopoiesis (EMH) | 2 | 5 | 7 |
| Oval cell hyperplasia | 2 | 2 | 4 |
| Kupffer cell hyperplasia | 2 | 2 | 4 |
| Biliary cystadenoma | 2 | 0 | 2 |
| Alveolar histiocytosis | 1 | 1 | 2 |
